# *Lukea* gen. nov. (Monodoreae-Annonaceae) with two new species of shrub from the forests of the Udzungwas, Tanzania & Kaya Ribe, Kenya

**DOI:** 10.1101/2021.05.14.444227

**Authors:** Martin Cheek, W.R. Quentin Luke, George Gosline

## Abstract

A new genus, *Lukea* Gosline & Cheek (Annonaceae), is erected for two new species to science, *Lukea quentinii* Cheek & Gosline from Kaya Ribe, S.E. Kenya, and *Lukea triciae* Cheek & Gosline from the Udzungwa Mts, Tanzania. *Lukea* is characterised by a flattened circular bowl-shaped receptacle-calyx with a corolla of three petals that give the buds and flowers a unique appearance in African Annonaceae. Both species are extremely rare shrubs of small surviving areas of lowland evergreen forest under threat of habitat degradation and destruction and are provisionally assessed as Critically Endangered and Endangered respectively using the IUCN 2012 standard. Both species are illustrated and mapped. Material of the two species had formerly been considered to be possibly *Uvariopsis* Engl. & Diels, and the genus *Lukea* is placed in the *Uvariopsis* clade of the Monodoreae (consisting of the African genera *Uvariodendron* (Engl. & Diels) R.E.Fries, *Uvariopsis, Mischogyne* Exell, *Dennettia* Bak.f., and *Monocyclanthus* Keay).

The clade is characterised by often conspicuous, finely reticulate quaternary nervation and incomplete or absent connective shields (in Annonaceae the connective shield is usually complete). Morphologically *Lukea* is distinct for its broad, turbinate, fleshy pedicel, a synapomorphy. It appears closest to the West African monotypic *Monocyclanthus*, sharing a trait unusual in the Annonaceae: the calyx in both genera forms a shallow bowl (calyx lobes are absent or vestigial), barely enclosing the base of the petals at anthesis, and persisting as a living, leathery disc at least until the fruit is mature. The placement of *Lukea* within the *Uvariopsis* clade is discussed.

## Introduction

In December 1953, *Semsei* 1520 (EA, K) was collected from Mtibwa Forest Reserve, Morogoro District, Tanzania. Sterile, it was initially tentatively identified as possibly a *Polyalthia* Blume, later as probably a *Uvariodendron* (Engl. & Diels) R.E.Fries, although this was doubted by Verdcourt. More than 40 years later Luke collected *Luke* 4703 (EA, K) also sterile, from Kaya Ribe in the coastal forests of Kenya. Recollected in Sept. 1997 (*Luke* & *Luke* 4740 (K) with flower buds and fruit, the latter two specimens were identified as “Annonaceae Genus Nov.” by Vollesen at Kew in 1998 and set aside for description. A further collection of the taxon from Kaya Ribe with open flowers and fruit was made in Jan. 1999 (*Luke* 5700, K). Two further collections were made of the Tanzanian taxon in 2003 and 2005, making flowers and fruit available for the first time (*Luke et al*. 9526, 11205, both K). Independently of Vollesen, both species were provisionally identified by Luke as *Uvariopsis* Engl. & Diels. Vollesen was not able to research the material further, being required to complete the Flora of Tropical East Africa account for Acanthaceae before his retirement (Darbyshire *et al*. 2010). The material of his new genus was passed to Gosline in 2017 together with other unresolved East African Annonaceae. Two of these, *Marshall* 1567 and *Cribb & Grey-Wilson* 10082, were placed in *Mischogyne* Exell using molecular methods and prompted a revision of that Afrotropical genus (Gosline *et al*. 2019). The new genus identified by Vollesen was placed by Gosline in Monodoreae in the region of *Uvariopsis* while noting the similarity of venation with *Mischogyne*. These two genera are placed in a clade together with *Uvariodendron* and *Monocyclanthus* (referred to here as the *Uvariopsis* clade) by Guo *et al*. (2017) and Gosline *et al*. (2019). The position remained uncertain until a molecular phylogenetic analysis of the whole Monodoreae tribe using a new set of nuclear markers (Couvreur *et al*. 2019) was undertaken by Dagallier & Couvreur (Dagallier *et al*. in prep.). This phylogenetic study was based on around 300 nuclear genes sequenced for almost all species of the tribe. Preliminary results indicated that the sequenced specimen *Luke* 11205 (type specimen of *Lukea triciae*) indeed clustered in Monodoreae, and more specifically in the informal “*Uvariopsis* clade” (which to date contains five genera *Dennettia, Mischogyne, Monocyclanthus, Uvariodendron* and *Uvariopsis* (Guo *et al*. 2017)). (The study also supports the resurrection of *Dennettia*, sunk into *Uvariopsis* by Schatz (Kenfack et al. 2003) treated here as a distinct genus.) The specimen was further recovered with maximum support as sister to *Mischogyne* (Dagallier, pers. comm.). However, both *Lukea* and *Mischogyne* showed important genetic differentiation (long branches separating both genera), confirming the new genus status as indicated by Vollesen. To date, no specimen of *L. quentinii* has been sequenced.

Twenty-seven genera are accepted in the Flora of Tropical East Africa account of Annonaceae, with 80 named species, and a further 11 which remained informally named (Verdcourt 1971). *Lukea* keys out in Verdcourt’s key to genera for flowering material, to couplet seven, since the carpels are free, and the hairs are simple. However, it fits neither arm of the couplet. While it has three valvate petals, it is not a tree, nor are the carpels 1-ovuled (*Enantia* Oliv., now *Annickia* Setten & Maas). Employing the key to fruiting material, it keys to couplet 54 because it has glabrous, non-tuberculate, several-seeded monocarps lacking longitudinal ribs with short stipes, seeds lacking pitted faces and far smaller than 2.5 cm long, leaves subglabrous. At this couplet it fails to fulfil both of the two statements since the fruiting pedicel (c. 1.5 mm long), neither exceeds 3 mm (leading to couplet 55) nor is the “fruit practically sessile” (the first arm: *Hexalobus* A.DC.). Further, in *Hexalobus* the petals are transversely plicate and united at the base, differing from those of *Lukea* which are unfolded and free.

In the fifty years since publication of the Flora of Tropical East Africa account (Verdcourt 1971), a further 26 additional new species have been described, bringing the cumulative total of formally named species of Annonaceae to 106 (Vollesen 1980; Verdcourt 1986; Verdcourt & Mwasumbi 1988; Johnson *et al*. 1999; Deroin & Luke 2005; Couvreur *et al*. 2006; Couvreur *et al*. 2009; Couvreur & Luke 2010; Marshall *et al*. 2016; Johnson *et al*. 2017; Gosline *et al*. 2019; Dagallier *et al*. 2021). It seems likely that additional species will continue to emerge for many years so long as taxonomists and botanical inventory work procede and natural habitat survives and is incompletely surveyed.

Here we formally describe the new genus as *Lukea* Gosline & Cheek including two new species *L. quentinii* Cheek & Gosline (Kenya) and *L. triciae* Cheek & Gosline (Tanzania). We compare *Lukea* morphologically with the genera in the *Uvariopsis* clade: *Uvariodendron, Uvariopsis, Mischogyne, Monocyclanthus* Keay and *Dennettia* Bak.f.

## Materials and methods

Herbarium citations follow Index Herbariorum (Thiers *et al*. 2020). Specimens were viewed at EA (using http://eaherbarium.museums.or.ke/eaherbarium/results accessed 28 April 2021), BR, K, MO (https://www.tropicos.org/home accessed 28 April 2021). All specimens cited have been seen by one or more of the authors.

Binomial authorities follow the International Plant Names Index (IPNI, 2020). The conservation assessment was made using the categories and criteria of IUCN (2012). Spirit preserved material was available of both species. Herbarium material was examined with a Leica Wild M8 dissecting binocular microscope fitted with an eyepiece graticule measuring in units of 0.025 mm at maximum magnification. The drawing was made with the same equipment using Leica 308700 camera lucida attachment. The flowers and fruits of herbarium specimens of the new species described were soaked in warm water to allow dissection. The terms and format of the description follow the conventions of Beentje & Cheek (2003) and Gosline *et al*. (2019). Georeferences for specimens lacking latitude and longitude were obtained using Google Earth (https://www.google.com/intl/en_uk/earth/versions/). The map was made using QGIS (https://www.qgis.org).

### Taxonomic Results

Morphological characters separating *Lukea* from other genera in *Uvariopsis* clade of Monodoreae are given in Table 1 below. *Lukea*, apart from the fleshy turbinate pedicel (a potential synapomorphy) has a combination of characters seen in no other genus.

**Table 1.**
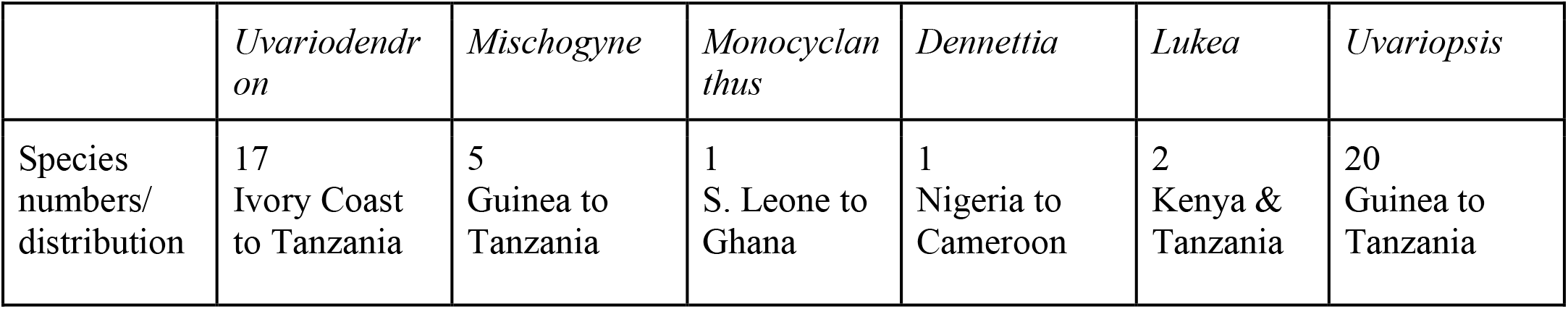
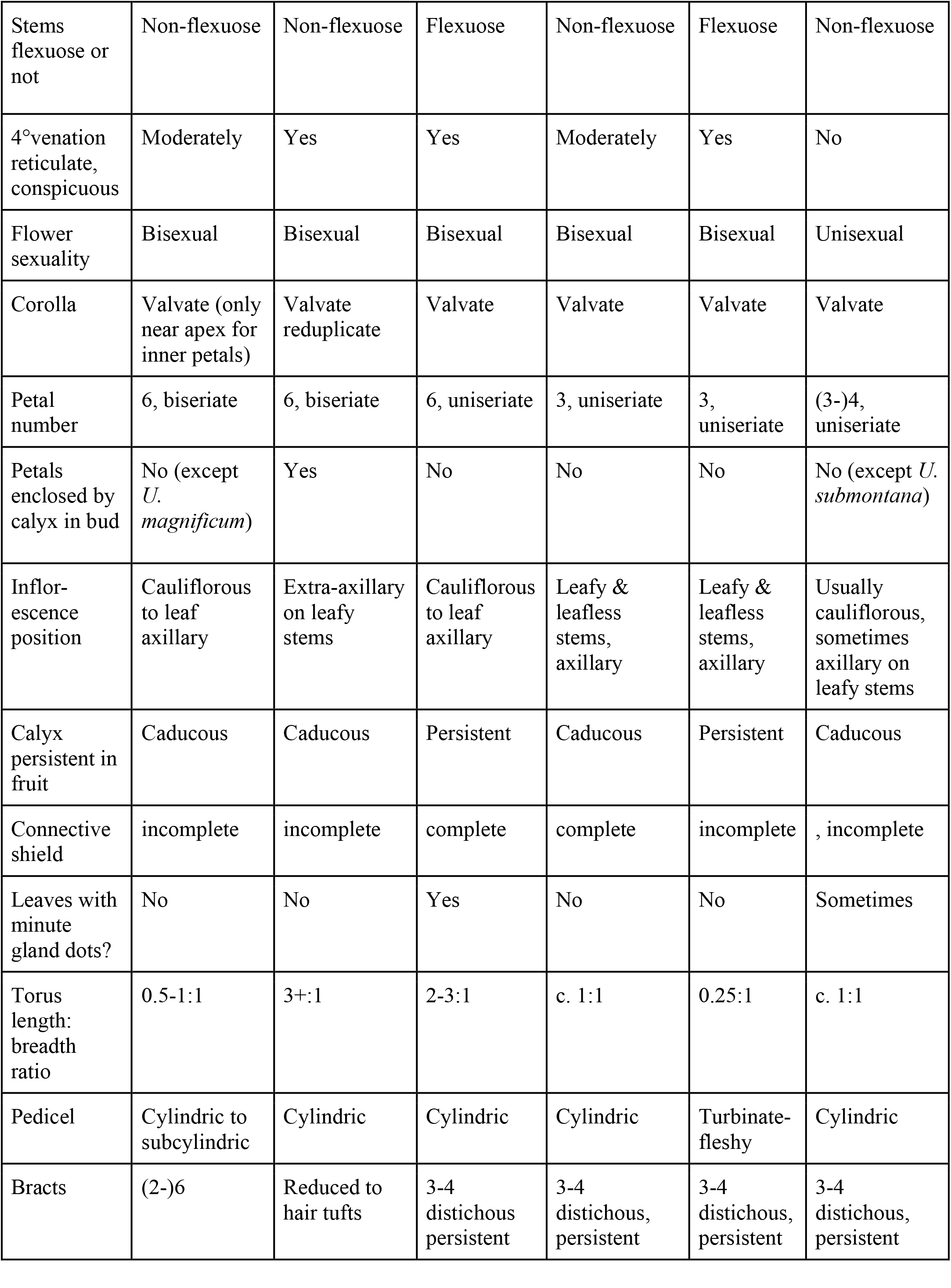
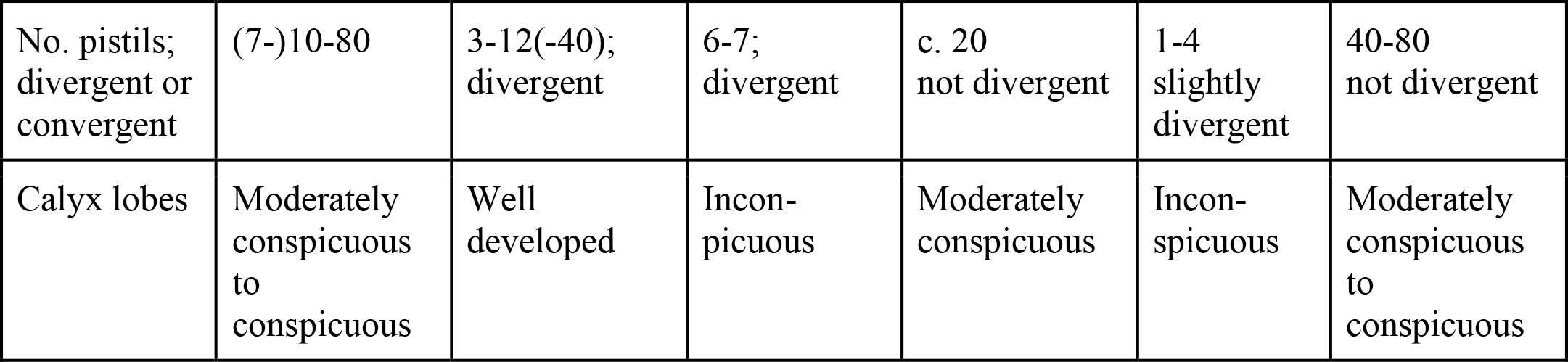
Diagnostic characters separating the genera of the *Uvariopsis clade: Uvariodendron, Mischogyne, Monocyclanthus, Dennettia, Lukea*, and *Uvariopsis*.

**Table 2.** Diagnostic features separating *Lukea quentinii* and *L. triciae*. Vegetative characters are based on those of reproductive (flowering and fruiting) stems.

|  | *Lukea quentinii* | *Lukea triciae* |
| --- | --- | --- |
| Indumentum of stems | Softly hairy, persistent, conspicuous, covering 10-20% of surface, hairs 0.1-0.2 mm long | Inconspicuous, soon glabrescent, hairs 0.05 mm long |
| Number of lateral nerves on each side of the midrib | 5-7 | 17-19 |
| Leaf blades shape and dimensions | Ovate or ovate-elliptic (4.5-)6.8-10.5 x (2.5-)4-6 cm | Oblong or elliptic-oblong (11.2-)13.9-21 x (4.4-)5.2-7.4(-8.1) cm |
| Petiole length | c. 3 mm | 5-9 mm |
| Width of petals | 5-6 mm | 8mm |
| Fruit | Cylindric, stipitate, constricted between the seeds | Ellipsoid, sessile, not constricted between the seeds |
| Number of pistils/flowers | 1-3 | 4 |
| Geography | Coastal S.E. Kenya (Kaya Ribe) | Tanzania (Morogoro Region) |

## Key to genera of the *Uvariopsis* clade

1. Quaternary nerves of leaf blades moderately or very conspicuous, forming a fine reticulum; petals & sepals 3-merous; flowers bisexual...................................................2
1. Quaternary nerves of leaf blades absent or inconspicuous; petals (4) and sepals (2) 2-merous; flowers unisexual (except *U. bisexualis*)...................................................***Uvariopsis***
2. Inflorescences extra-axillary; bracts reduced to hair-tufts; calyx lobes covering corolla in bud; petals biseriate, strongly reflexed at anthesis...................................................***Mischogyne***
2. Inflorescence axillary (or sometimes cauliflorous); bracts not reduced to hair tufts but fully developed, 3 -4, distichous; calyx lobes if present, not covering corolla in bud; petals uniseriate, if reflexed, incompletely and slightly so...................................................3
3. Stems non-flexuose; calyx caducous; pistils 7-80, calyx lobes moderately conspicuous ....................................................................4
3. Stems flexuose; calyx persistent in fruit; pistils 1-7, calyx lobes absentabsent....................................................................5
4. Corolla biseriate................................................... ***Uvariodendron***
4. Corolla uniseriate ................................................... ***Dennettia***
5. Petals 6; pedicel cylindric; leaves with translucid gland dots, *citrus* scented (crushed); torus 2-3:1; pistils 6-7...................................................***Monocyclanthus***
5. Petals 3; pedicel turbinate; leaves without translucid gland dots, not citrus scented; torus 0.25:1; pistils 1-4...................................................***Lukea***

### *Lukea Gosline & Cheek* gen. nov

Type species: *Lukea quentinii* Gosline & Cheek

#### Hermaphrodite evergreen shrubs

Leafy stems distichous, producing flushes of 2 – 4 leaves per season terminating in a dormant bud; bud-scales orbicular-ovate; stems terete, often flexuose (zig-zag), first (current) season’s growth drying green, lacking ridges and lenticels, indumentum sparse, hairs of 1 or 2 size-classes, simple, translucent, erect; axillary buds a dome-shaped, of numerous filamentous, densely hairy scales. Second season’s epidermis greyish black, with low, sinuous longitudinal ridges; lenticels inconspicuous, concolorous, raised, longitudinally elliptic, with a median longitudinal fissure; glabrescent.

*Leaves* persisting for two seasons or more, stage-dependent heteromorphic (*L. quentinii*): those of sterile (non-reproductive) stems longer and larger than those of reproductive stems. *Leaf-blades* drying green, lacking translucid gland dots, subglossy or glossy above and matt below, ovate, elliptic or elliptic-oblong, rarely slightly oblanceolate, acumen short, broadly triangular, base rounded; midrib cylindric on the abaxial surface, visible usually only as a slit on the upper surface. Secondary nerves 6 – 19 on each side of the midrib, brochidodromous, arising at c. 60 – 80° from the midrib, straight, forking at c. 1 cm from the margin and uniting with the secondary nerves above and below to form an angular inframarginal nerve, connected by lower order nerves to two other parallel and less distinct, inframarginal nerves closer to the thickened, revolute margin. Tertiary and quaternary nerves also raised, as prominent as the secondary nerves, and forming a conspicuous reticulum of isodiametric cells on both adaxial and abaxial surfaces, cell (areolae) diameter c. 1 mm. Indumentum absent or sparse and inconspicuous, along the abaxial midrib and margin, hairs simple, appressed, white. Petioles slightly twisted, dark green drying dull or bright orange, terete but the adaxial surface with a median slit, narrower in width at base and apex, transversely ribbed, glabrous or sparsely hairy.

*Inflorescences*, axillary, scattered sparsely along leafy or leafless nodes of stems 2 – 4 seasons old, 1-flowered, often (*Lukea quentinii*) with a second inflorescence arising sequentially from the base of the first. Peduncle and rachis together very short, hairs scattered, simple, dark brown, appressed. Bracts 3 – 4, distichous, green, persistent, shortly ovate-elliptic, apex acute or mucronate, base sheathing or not, increasing in size towards the pedicel, hairs sparsely scattered. Pedicel fleshy-leathery, yellow-green, obconical, widening steadily from base to apex, the hairs as in the bracts. *Flowers*, bisexual. Calyx-receptacular tube, thick, leathery, externally warty, broad and short, shallow bowl-shaped, partly enclosing corolla in bud, indumentum on outer surface of scattered, dark brown, simple hairs, inner surface glabrous. Calyx lobes absent or vestigial, 3, alternating with the petals, triangular, minute, inserted at the edge of the calyx-receptacular tube aperture, centripetal (directed towards the floral axis), persistent (if developed), indumentum as receptacle but denser. Petals 3, alternating with sepals, green, valvate, thick and fleshy, shallowly concave, appearing nearly flat (horizontal) in bud (petal axis nearly perpendicular to that of floral axis), at anthesis elevated from the horizontal by 40 – 70°, triangular, apex acute, base truncate-retuse, sides square, outer surface densely covered in appressed sericeous golden hairs, inner surface glabrous. Torus (raised surface of receptacle on which are inserted the stamens and carpels), domed, wider than long. Stamens c. 150 – 180, spirally inserted, arranged in 8 – 12 ranks. Stamens latrorse, basifixed, erect, linear, connective and thecae nearly the length of the stamen, connective terminal extension globose, black, forming an incomplete cap over the anther cells. Pollen in tetrads. Petals and stamens falling after anthesis leaving the calyx-receptacle cup and carpels exposed, protected in the calyx-receptacular tube. Carpels (1 –)2 – 4, brown, not diverging from each other, longer than stamens, subcylindrical, densely brown pubescent, the hairs patent. Stigma sessile, horseshoe shaped-crescentic, adaxial surface glabrous, outer surface with minute white patent hairs. Ovules numerous, biseriate, parietal. *Fruit* monocarps 1(– 2) (numbers per fruit unknown in *L. triciae*), cylindric or narrow ellipsoid, constricted or not between the seeds, yellow when mature, pericarp leathery, glossy, glabrous; shortly stipitate or sessile, glabrous or glabrescent; apex rounded, obscurely bifurcate. Fruiting pedicels slightly accrescent. Calyx-receptacle and bracts persistent, green, not accrescent. *Seeds* 3 – 12 per monocarp, orange-brown discoid, or pale-yellow oblong-elliptic and widest near hilum, hilum large, elliptic, with a ragged, crater-like rim, raphe slightly raised, girdling the seed, testa bony, lacking pitting or sculpture; endosperm conspicuously ruminate in longitudinal section, embryo flat, minute,

##### RECOGNITION

Similar to *Monocyclanthus* Keay in the flexuose stems, conspicuous fine quaternary reticulate venation, axillary, 1-flowered inflorescences (sometimes cauliflorous in *Monocyclanthus*), bisexual flowers, and calyx-receptacle with only rudimentary (or no) lobes, persistent and disc-like in fruit; differing in leaves lacking translucid gland dots and *Citrus* scent, torus broader than long (vs twice or more as long as broad), petals 3 (vs 6), the turbinate fleshy pedicel (vs cylindric), pistils 1 – 4 per flower (vs 6 – 7).

##### DISTRIBUTION

S.E. Kenya, Kilifi County and Tanzania, Morogoro region. The distribution of *Lukea*, with one species in the Eastern Arc Mts and another in the coastal forests of Kenya is seen in several other genera, such as *Ancistrocladus* Wall. with *A. tanzaniensis* Cheek *& Frim*.*-Møll. in the Udzungwas and A. robertsoniorum* J.Léonard in the Kenyan coastal forests (Cheek *et al*. 2000, Cheek 2000), also in the genus *Afrothismia* with *A. mhoroana* Cheek in the Ulugurus and *A. baerae* Cheek in Kenyan coastal forests (Cheek 2004a; Cheek 2006; Cheek & Jannerup 2006). Numerous other taxa are restricted to the Eastern Arc Mts of Tanzania and the Kenyan Coastal Forests, which together are referred to as EACF (see discussion).

**Map 1.**
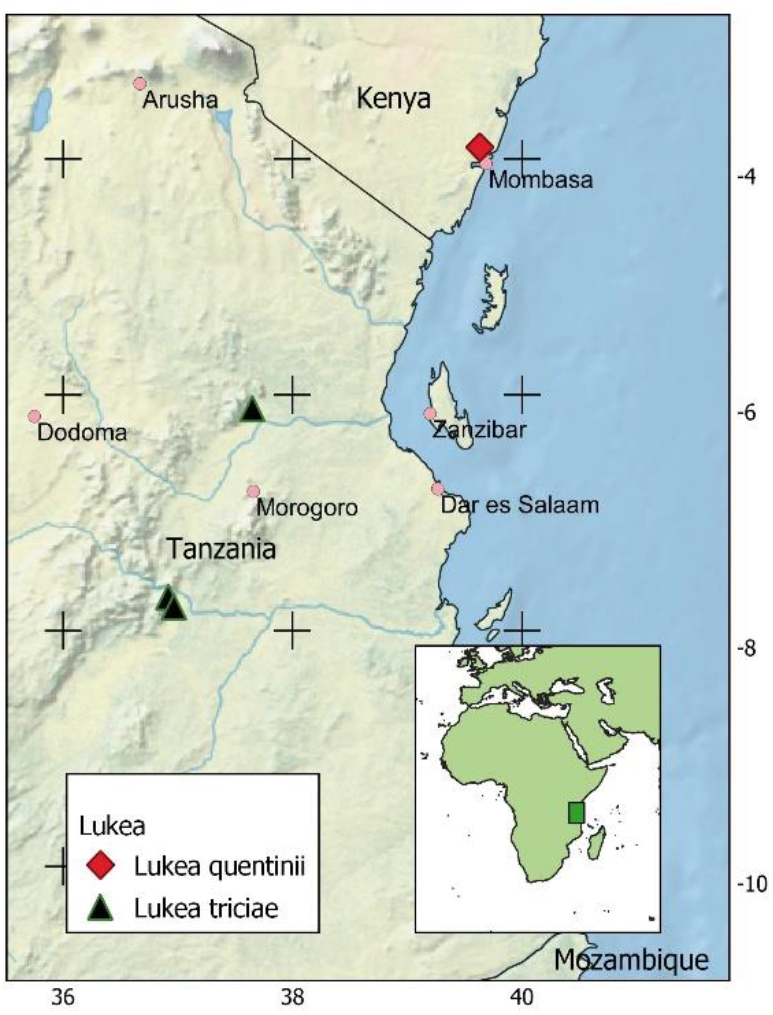
Global distributions of *Lukea quentinii* and *L. triciae*.

##### HABITAT

Evergreen lowland forest remnants, c. 50 – 300 m elevation.

##### CONSERVATION STATUS

both species are globally threatened (see below under the species accounts).

##### ETYMOLOGY

Named for the noted Kenyan botanist, Research Associate of the East African herbarium (EA), William Richard Quentin Luke, better known as Quentin Luke (1952 –). He is the leading specialist on the plants of the Kenyan coastal forests, and has discovered numerous new species of plants especially in Kenya and Tanzania, such as the incredible spectacular Tanzanian tree acanth *Barleria mirabilis* I.Darbysh. & Q.Luke (Darbyshire & Luke 2016). He has also brought to light species from across Africa e.g., in Democratic Republic of Congo: *Keetia namoyae* O. Lachenaud & Q. Luke (Lachenaud *et al*. 2017) and from Mali and Guinea the only endemic African *Calophyllum, C. africanum* Cheek & Q. Luke (Cheek & Luke 2016; Couch *et al*. 2019). He has also collected and described many other novel Annonaceae from Tanzania and Kenya (see introduction). Ten species are named for him, including the Tanzanian species *Cola quentinii* Cheek (Cheek & Dorr 2007) and *Cola lukei* Cheek (Cheek 2002). He is the principal collector of all known fertile material of the two species comprising the genus *Lukea*. For further biographical and bibliographical information see (Polhill & Polhill 2015: 276 – 277).

##### NOTES

Local names and uses of the species are unknown and may not exist since both species are so rare. The pollination biology is also unknown, but we conjecture that beetles may be involved as is usual in Annonaceae in Africa since the petals remain largely closed at anthesis forming a pollination chamber around the sexual parts (Gottsberger *et al*. 2011; Gottsberger 2012). Whether thermogenesis occurs as in *Uvariodendron* (Gottsberger *et al*. 2011) remains to be discovered.

##### Placement of *Lukea* in the *Uvariopsis* clade

The *Uvariopsis* clade of the Monodoreae (consisting of the African genera *Uvariodendron* (Engl. & Diels) R.E.Fries, *Uvariopsis* Engl. & Diels, *Mischogyne* Exell, *Dennettia* Bak.f., and *Monocyclanthus* Keay) is characterised (Table 1) by a tendency (present in most but not all genera of the clade) to conspicuous, finely reticulate quaternary nervation in the leaf-blades (usually absent or inconspicuous in Annonaceae) and incomplete or absent connective shields (in Annonaceae the connective shield is usually complete and is considered plesiomorphic). Placement of *Lukea* with this group is supported by possession of both these characters. The two characters are not seen in all genera e.g., in *Uvariopsis* the fine reticulation is not conspicuous, and a connective shield is present in *Monocyclanthus* and *Dennettia*. The last genus has been sunk into *Uvariopsis* (Kenfack *et al*. 2003) but is maintained as a separate genus here since it differs in having a connective shield, and in combining two characters (trimerous perianth and bisexual flowers) otherwise seen separately only in two atypical species of *Uvariopsis* (*Uvariopsis congolana* (De Wild.) R.E.Fr., and *U. bisexualis* Verdc., respectively). Despite *Lukea* being placed in the *Uvariopsis* clade as sister to *Mischogyne* (Dagallier in prep.), it lacks the numerous diagnostic features of that genus within the clade, such as the extra-axillary inflorescences, the calyx-sheathed corolla in bud, the strongly reflexed corolla with exserted pistils and the extremely long torus (length: breadth 3+:1) (Gosline *et al*. 2019). Similarities are the reticulate venation, the stamens without connective shield, and the ellipsoid monocarps in the fruit.

Morphologically the new genus appears closest to the West African monotypic *Monocyclanthus*, sharing a trait unusual in the Annonaceae: the calyx in both genera forms a shallow bowl (calyx lobes are absent or vestigial), barely enclosing even the base of the petals at anthesis. The bowl persists as a live, leathery disc at least until the fruit is mature. A similar structure is found in *Uvariodendron schmidtii* Q.Luke, Dalgallier & Couvreur (Dagallier *et al*. 2021) in which the three calyx lobes are distinct, but appear almost completely connate (the individual lobes can still be clearly discerned) forming a shallow saucer at anthesis, however this structure falls post-anthesis (Dagallier *et al*. 2021) unlike in *Lukea*. The ancestor of *Lukea* may have had a similar calyx, which may have evolved by developing greater concavity, more complete loss of the calyx lobes, and persistence in the fruit. The occasional presence of vestigial inner petals in *Lukea* also suggest a link with *Mischogyne* and *Uvariodendron* which alone in the clade have regular inner petals. The function of the calyx-like structure, which may derive more from the enlarged receptacle than the calyx (suggested by the lack of evident calyx lobe connation) apart from protecting the flower in early bud, is unknown. It may be that the same structure in *Monocyclanthus* arose independently, and that the close similarity with *Lukea* does not indicate a sister relationship. *Monocyclanthus* and *Lukea* also share flexuose leafy twigs (seen intermittently in genera of other tribes e.g. *Mwasumbia* Couvreur & D.M. Johnson), and an unusually low number of pistils per flower (1 – 4 in *Lukea* and 6 – 7 in *Monocyclanthus*).

## Key to the species of *Lukea*

Stems of leafy reproductive shoots thinly but softly white hairy, hairs patent, 0.1 – 0.2 mm long; leaves ovate-elliptic, (4.5 –)6.8 – 10.5 x (2.5 –)4 – 6 cm, lateral nerves 5 – 7 on each side of the midrib; petals 5 – 6 mm wide; fruit stipitate, cylindric, constricted between the seeds. Coastal Kenya (Kaya Ribe) **……………………………….…**..**1. *Lukea quentinii***

Stems of leafy reproductive shoots glabrescent, hairs inconspicuous, 0.05 mm long; leaves ovate-elliptic, (11.2 –)13.9 – 21 x (4.4 –)5.2 – 7.4(– 8.1) cm, lateral nerves 17 – 19 on each side of the midrib; petals 7 – 9 mm wide; fruit sessile, narrowly ellipsoid, not constricted between the seeds. Tanzania (Morogoro District) ………………………….. **2.*Lukea triciae***

### 1. Lukea quentinii

Gosline & Cheek *sp. nov*. Type: Kenya, Kilifi County, Kaya Ribe, Mleji River, 03°54’ S, 39°38’ E, fl., fr. 16 Jan. 1999, *Q. Luke* 5700 (holotype K000593320; isotypes EA000008943, MO, US).

#### *Hermaphrodite evergreen shrub* c 3 m tall

Leafy stems producing flushes of 3 – 4 leaves per season terminating in a dormant bud; bud-scales c. 4 – 7, orbicular 1.5 × 1.5 mm; stems terete, 2 – 3 mm diam. often flexuose (zig-zag), internodes 2 – 4.2 cm long, first (current) season’s growth drying green, lacking ridges and lenticels, indumentum moderately dense, covering 10 – 20% of the surface, hairs simple, white, erect, 0.1 – 0.2 mm long, intermixed with a few longer hairs, persisting for six internodes or more from the stem apex. Second season’s growth greyish black, with low, sinuous longitudinal ridges; lenticels inconspicuous, concolorous, raised, longitudinally elliptic, 0.6 – 1.5 × 0.5 mm, with a median longitudinal fissure; glabrescent. *Leaves* persisting for two seasons or more, blades drying pale green, lacking glands, thickly papery, subglossy above and matt below, those of sterile (non-reproductive) stems (*Luke* 4703, K) longer than those of reproductive stems, elliptic or elliptic-oblong, rarely slightly oblanceolate, 11 – 15 × 5.5 – 7.8 cm, acumen short, broadly triangular, c. 0.4 × 0.4 cm, base rounded, minutely and abruptly cordate at the midrib; midrib cylindrical, 1 – 1.2 mm diam. on the abaxial surface, visible only as a slit on the upper surface, densely patent hairy, glabrescent. Secondary nerves 7 – 8 on each side of the midrib, brochidodromous, arising at c. 80° from the midrib, straight, forking at c. 1 cm from the margin and uniting with the secondary nerves above and below to form an angular inframarginal nerve, connected by lower order nerves to two other, less distinct, inframarginal nerves closer to the thickened, revolute margin. Tertiary and quaternary nerves also raised, as prominent as the secondary nerves, and forming a conspicuous reticulum on both adaxial and abaxial surfaces. Indumentum sparse and inconspicuous, along the abaxial midrib and abaxial blade towards the margin, hairs simple, appressed, white, 0.1(–0.3) mm long. Petioles dull or bright orange, terete but the adaxial surface with a median slit, (4.5 –)6 mm long, (1.5 –)2.5 mm wide, narrower in width at base and apex, transversely ribbed, glabrous. Leaves of reproductive stems (*Luke* 4740, 5700, K) similar to but smaller and differently shaped to those of the sterile stems, ovate or ovate-elliptic, (4.5 –)6.8 – 10.5 x (2.5 –)4 – 6 cm, lateral nerves 5 – 7(– 8) on each side of the midrib. Petioles cylindric 3 × 1.3 mm, sparsely covered (5 – 10% of surface) with patent white hairs 0.3 mm long. Inflorescences, axillary, scattered sparsely along leafy or leafless nodes of stems 2 – 4 seasons old, 1-flowered, often with a second inflorescence arising sequentially from the base of the first. Peduncle and rhachis together 0.5 – 1 × 0.5 – 1 mm at anthesis, hairs dense becoming scattered with age, simple, dark brown or bronze, 0.2 – 0.5 mm long.

Bracts 3 – 4, distichous, green, persistent, shortly ovate-elliptic, 1.2 – 1.5 × 1.2 – 1.5 mm apex acute or mucronate, base sheathing, increasing in size towards the pedicel, hairs scattered, red-brown 0.1 – 0.2 mm long. Pedicel obconical, fleshy-leathery, yellow-green, widening steadily from base to apex, the base 1.5 mm diam., apex 3.5 mm diam., hairs as bracts. *Flowers*, bisexual. Calyx-receptacular tube mostly enclosing corolla in bud, bowl-shaped, 2 mm long, 8 mm wide, distal aperture orbicular, 6 – 8 mm diam. at anthesis, indumentum on outer surface scattered, dark brown, simple 0.25 – 0.3 mm long, inner surface glabrous. Calyx absent or vestigial, lobes 3, small triangular 0.1 – 0.2 × 0.5 mm, inserted at the edge of the receptacular tube distal aperture, centripetal, indumentum as receptacle. Petals 3, alternating with sepals, green, valvate, thick and fleshy, slightly concave, appearing nearly flat (horizontal) in bud (petal axis nearly perpendicular to that of floral axis), at anthesis elevated from the horizontal by 40 – 50°, triangular, 5 × 5 – 6 mm, apex acute, base (when detached) truncate-retuse, sides square, 1.5 mm thick along the midline, outer surface densely covered in appressed sericeous golden hairs. Vestigial inner petal sometimes present, oblanceolate to spatulate, 2.5 × 0.75 mm, apex rounded, glabrous. Torus (raised surface of receptacle on which are inserted the stamens and carpels), domed, wider than long, 0.5 – 0.75 × 2 mm. Stamens c. 150, spirally inserted, arranged in 8 – 10 ranks. Stamens latrorse, basifixed, erect, linear 1.4 – 1.5 × 0.25 mm, filament 0.1 mm long, connective and thecae nearly the length of the stamen, connective terminal extension globose, 0.1 mm long, black, forming an incomplete cap over the anther cells. Petals and stamens dropping after anthesis leaving the calyx-receptacle cup and carpels exposed. Carpels (1 –)2(–3) brown, longer than stamens, subcylindrical, 2.5 × 0.5 mm, densely brown pubescent, the hairs patent, 0.05 – 0.1 mm long. Stigma yellowish white, sessile, horseshoe shaped-crescentic, 0.8 × 0.6 mm, adaxial surface glabrous, outer surface with minute white patent hairs 0.05 mm long. *Fruit* monocarps 1(– 2), cylindrical, 1.8 – 2.3(–3.5) × 1.1 – 1.3(– 1.5) cm, constricted c. 1 mm between the (3 –)4 – 6 seeds, yellow, pericarp leathery, glossy, glabrous apart from a few simple hairs 0.2 mm long; stipe 1 – 1.5 × 2 mm, glabrous; apex rounded, minutely bifurcate. Fruiting pedicels slightly accrescent, 1.5 mm long. Calyx-receptacle and bracts persistent in fruit, green, not accrescent. *Seeds* 3 – 6 per monocarp, orange-brown, discoid, 10 – 11 mm diam. × 4 – 6 mm thick, margin rounded, testa 0.25 mm thick, raphal cavities 1 – 1.2 mm diam.; hilum elliptic 4 – 6 × 1 – 2 mm, rim crater-like, 0.5 mm high, verrucose; endosperm ruminate, embryo transversely elliptic in transverse section, 4.5 × 2.5 mm. Fig. 1.

**Fig. 1.**
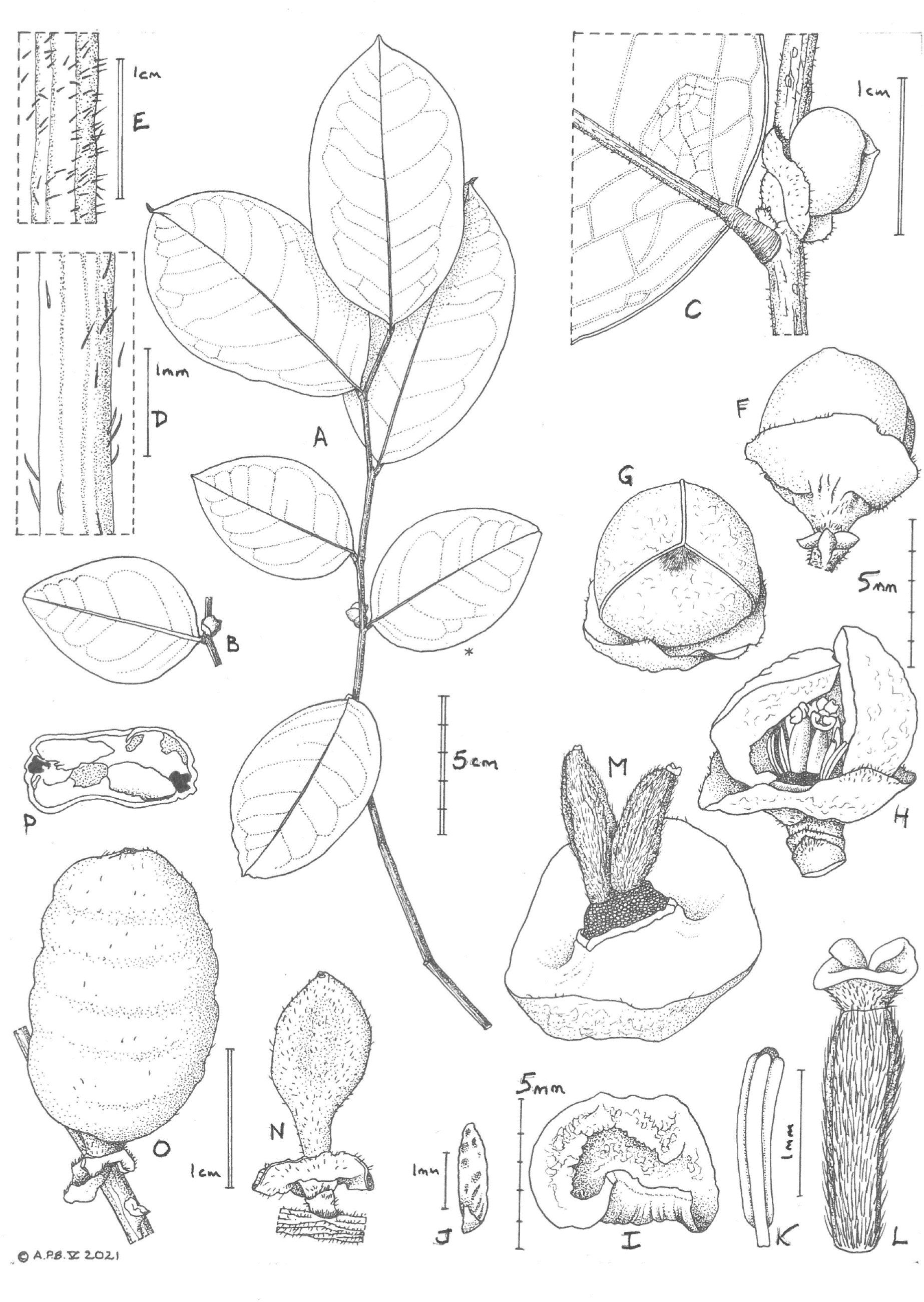
*Lukea quentinii* **A** habit, leafy, flowering stem, leaves showing adaxial surface**; B** habit, flowering node, showing abaxial surface of blade; **C** close-up of B; **D** hairs on abaxial midrib; **E** hairs on distal stem internode; **F** flower bud, lateral view; **G** flower bud, plan view; **H** flower with petal removed to show stamens and pistils; **I** petal, abaxial surface; **J** vestigial inner petal; **K** stamen, showing lateral pollen thecae and incomplete connective cap; **L** pistil, side view; **M** flower post anthesis/young fruits; **N** immature fruit with persistent calyx; **O** mature fruit; **P** seed, transverse section. **A-E, G-M, P** from *Luke et al*. 5700; **F, N, O** from *Luke et al*. 4740. All drawn by ANDREW BROWN.

##### RECOGNITION

Similar to *Lukea triciae* Cheek & Gosline, distinct in the leaves of reproductive stems smaller ((4.5 –)6.8 – 10.5 × (2.5 –)4 – 6 cm long), ovate or ovate-elliptic with lateral nerves 5 – 7(– 8) on each side of the midrib (versus (11.2 –)13.9 – 21 × (4.4 –)5.2 – 7.4(– 8.1) cm), oblong, lateral nerves 17 – 19 on each side) and in the smaller flowers (petals 5 – 6 mm wide versus 8 mm wide), carpels (1 –)2(– 3) (versus 4); fruits constricted between the seeds (versus not constricted between the seeds).

##### DISTRIBUTION

S.E. Kenya (Map 1). Endemic to the Kaya Ribe forest, Kilifi County.

##### SPECIMENS EXAMINED

KENYA. Kilifi County, Kaya Ribe, 0354 S, 3938 E, 15 July 1997, *Q. Luke* 4703 (EA000008944, K000593319); ibid, Mleji River fl. buds, fr. 16 Sept. 1997 *Q. Luke & P. Luke* 4740 (EA, K000593318, MO n.v.); ibid., fl., fr. 16 Jan. 1999, *Q. Luke* 5700 (K000593320 holo.; iso: EA000008943, MO, US n.v.).

##### HABITAT

Evergreen to semi-deciduous coastal Kaya forest along stream. Associated species are *Encephalartos hildebrandtii, Asteranthe* sp. nov., *Mkilua fragrans, Uvariodendron kirkii, Xylopia holtzii, Gyrocarpus americanus, Rinorea* spp., *Barringtonia racemosa, Combretum schumannii, Terminalia sambesiaca, Garcinia* spp., *Cola* spp., *Sterculia* spp., *Rhodognaphalon schumannianum, Euphorbia wakefieldii, Ricinodendron heudelotii, Aristogeitonia monophylla, Afzelia quanzensis, Bauhinia mombassae, Caesalpinia insolita, Cynometra* spp., *Dialium* spp., *Hymenaea verrucosa, Julbernardia magnistipulata, Newtonia paucijuga, Parkia filicoidea, Cordyla africana, Erythrina sacleuxii, Millettia* spp., *Celtis* spp., *Antiaris toxicaria, Ficus* spp., *Milicia excelsa, Trichilia emetica, Lecaniodiscus fraxinifolius, Lannea welwitschii, Sorindeia madagascariensis, Diospyros* spp., *Manilkara* spp., *Synsepalum* spp., *Breonadia salicina, Premna* spp., *Borassus aethiopum, Pandanus rabaiensis* (Luke unpublished data). The forest is on a hill c. 90 m high that rises from a plain at 40 – 50 m alt.

##### CONSERVATION STATUS

Kaya Ribe, the sole known location for *Lukea quentinii*, is one of the Kenyan coastal Kaya forests. It is a traditional sacred forest of the Mjikenda and in particular for the WaRibe tribe. It was gazetted as a National Monument (NM) in 1992 under the National Museums & Monuments Act. This has afforded it some protection but it still suffers from tree cutting and encroachment. It has also been listed with other Kayas as a World Heritage Site under UNESCO. From CNES/Airbus imagery dated 29 June 2020 available through Google Earth (viewed 8 May 2021, see 3.900 S 39.633E), the forest looks in reasonable condition although there appear to be several gaps, possibly from the threats mentioned above. In particular, since 2011 imagery shows that forest along the stream, which runs for c. 1 km through the forest, bisecting it, appears to have reduced in coverage. The forested stream edge is the main, possible the only, habitat of *Lukea quentini*, The forest is about 1 km diam., with an area of about 3.14 km^2^, equating to both the area of occupancy and extent of occurrence of the species. The forest is surrounded by agricultural fields. Numerous other plant species of the Kenyan coastal forests are already Red Listed, e.g. *Cola pseudoclavata* Cheek, *C. octolobioides* Brenan (Cheek & Lawrence 2019; Luke *et al*. 2018 respectively). We here provisionally assess *Lukea quentinii* as Critically Endangered, CR B1+2ab(iii) given the metrics and threats stated above.

##### PHENOLOGY

sterile (lacking flowers and fruit) in July, flower buds present in Sept., open flowers, ripe and immature fruit in Jan.

##### ETYMOLOGY

named for Quentin Luke, as for the genus (see above).

##### VERNACULAR NAME & USES

None are recorded.

##### NOTES

The first known collected specimen of *Lukea quentinii* (*Q. Luke* 4703, K) is sterile. It has larger and longer leaves than the other two, fertile collections, and so we deduce that these leaves may represent a juvenile, or at least, a non-reproductive stage.

This species is extremely rare and localised. It represents an addition to the Flora of the Kenyan coastal forests, not being listed in Ngumbau *et al*. (2020).

### 2. Lukea triciae

Gosline & Cheek *sp. nov*. Type: Tanzania, Morogoro District, Udzungwa Mts National Park, Msolwa - Camp 244, 7° 43’ S, 36° 55’ E, 350 m alt., fl. 23 Oct. 2005, *Luke WRQ, Mwangoka, Festo* 11205 (holo. K000593317); iso. EA000008947, MO, NHR) (Fig. 2 & 3).

**Fig. 2.**
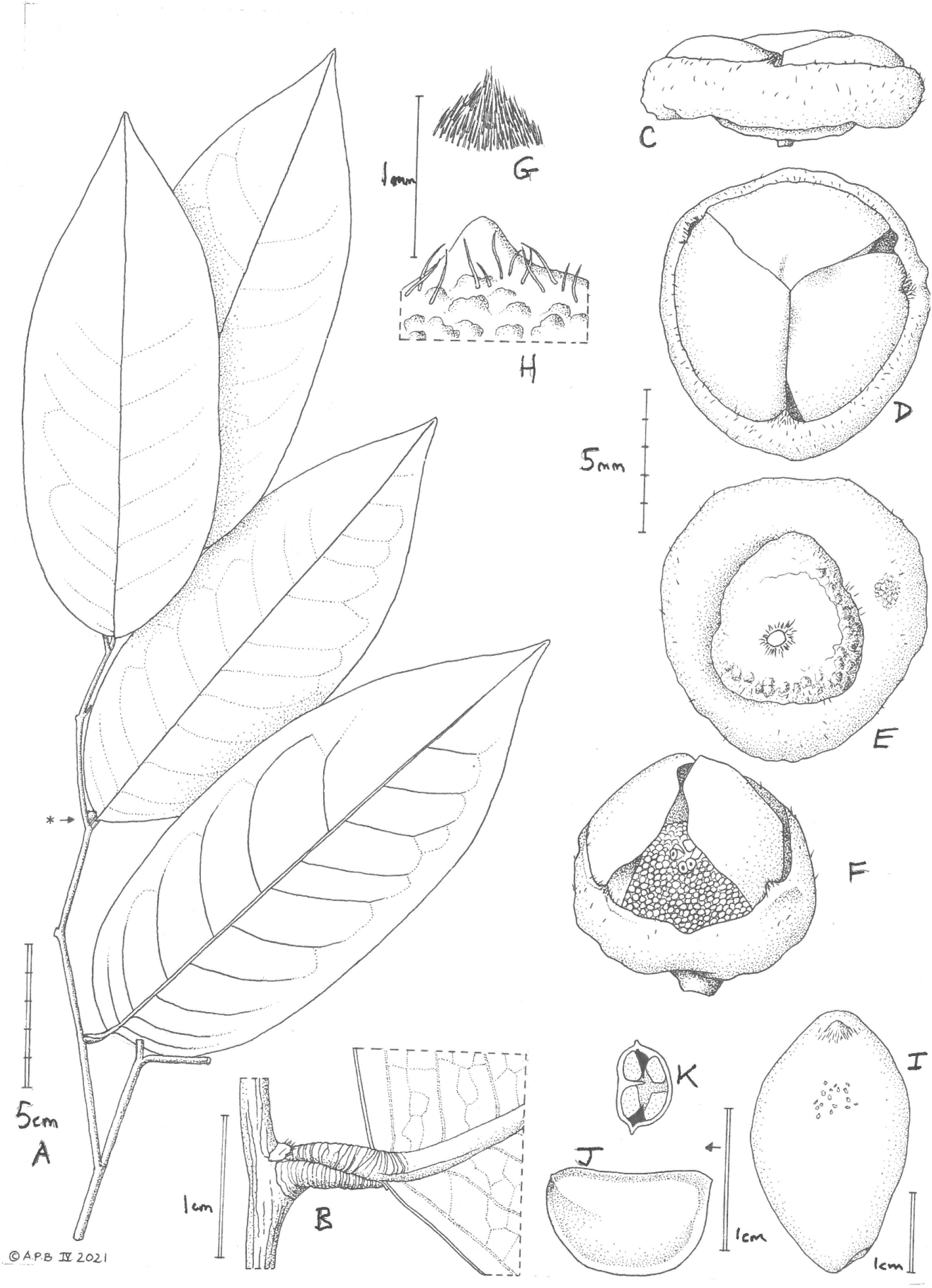
*Lukea triciae* **A** habit, leafy, flowering stem, leaves showing adaxial and (most distal leaf) abaxial surfaces; **B** detail of node showing abaxial leaf surface; **C** flower bud, side view; **D** flower bud, plan view; **E** flower bud, view from pedicel showing expanded calyx and the verrucate, turbinate, fleshy pedicel; **F** flower, petal removed to show stamens and pistils on torus; **G** hairs on outer surface of petals; **H** calyx, showing verrucate surface, indumentum, and a minute vestigial sepal lobe; **I** fruit, reconstructed; **J** seed, side view; **K** seed, transverse section. **A-H** from *Luke et al*. 11205; **I-K** from *Luke et al*. 9526; All drawn by ANDREW BROWN.

**Fig. 3.**
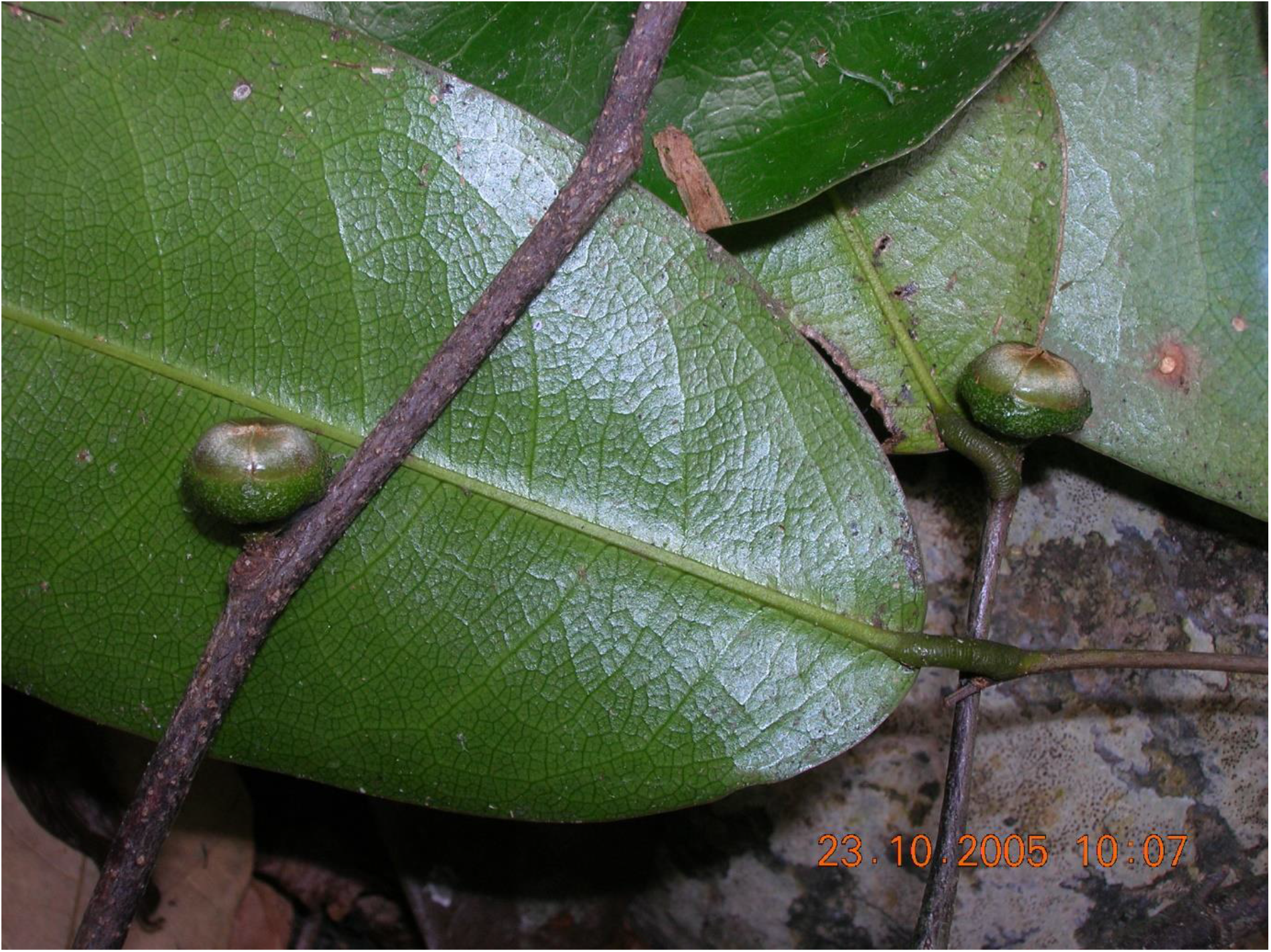
*Lukea triciae*. Flowering, leafy stem, from *Luke et al*. 11205. Photo by W.R.Q. Luke.

#### *Hermaphrodite evergreen shrub* or small tree 2 – 3 m tall

Leafy stems producing flushes of 2 – 3 leaves per season, terminating in a dormant bud; bud-scales 6 – 8, ovate 1.5 × 1.5 mm; stems terete, 2 – 3 mm diam. often flexuose (zig-zag), internodes 1.7 – 4 cm long, first (current) season’s growth drying green, lacking ridges and lenticels, indumentum glabrescent, inconspicuous, very sparse, simple, translucent, erect, >0.05mm long. Second season’s growth greyish black, with low, sinuous longitudinal ridges; lenticels inconspicuous, concolorous, raised, longitudinally elliptic, glabrescent. *Leaves* persisting for two seasons or more, blades drying coriaceous, elliptic-oblong, (11.2 –)13.9 – 21 × (4.4 –)5.2 – 7.4(– 8.1) cm, acumen c. 0.8 – 1.4 cm, base rounded; midrib cylindrical, 1 – 1.2 mm diam. on the abaxial surface, visible only as a slit on the upper surface. Secondary nerves 17 – 19 on each side of the midrib, brochidodromous, arising at c. 70 – 80° from the midrib, arching upwards, forking at 1 – 2 cm from the margin and uniting with the secondary nerves above and below to form an angular inframarginal nerve, connected by lower order nerves to two other, less distinct, inframarginal nerves closer to the thickened, revolute margin. Tertiary and quaternary nerves also raised, as prominent as the secondary nerves, and forming a conspicuous reticulum on both adaxial and abaxial surfaces. Indumentum sparse and inconspicuous, glabrescent. Petioles dull or bright orange, terete but the adaxial surface with a median slit, 5 – 9 mm long, (1.5 –)2.5 mm wide, narrower in width at base and apex, transversely ribbed, glabrous. *Inflorescences*, axillary, scattered sparsely along leafy or leafless nodes of stems 2 seasons old, 1-flowered. Peduncle and rhachis together c.1 mm long at anthesis, hairs sparse, simple. Bracts 3 – 4, distichous, persistent, shortly ovate, c.1.5 × c.1.5 mm apex rounded, base not-sheathing, increasing in size towards the pedicel, hairs scattered, red-brown 0.1 – 0.2 mm long. Pedicel obconical, fleshy-leathery, yellow-green, widening steadily from base to apex, the 1.5 – 3.5 mm long, base 1.5 mm diam., apex 5 – 6 mm diam., glossy, subglabrous. *Flowers*, bisexual, known only from slightly pre-anthetic material. Calyx-receptacular tube mostly enclosing corolla in bud, bowl-shaped, 2 mm long, 8 mm wide, distal aperture orbicular, 6 – 8 mm diam. at anthesis, indumentum on outer surface scattered, dark brown, simple 0.25 – 0.3 mm long, inner surface glabrous. Calyx vestigial, lobes 3, triangular, 0.5 × 1 mm, hairs moniliform, simple, 0.25 – 0.3 mm long. Petals 3, valvate, thick and fleshy, slightly concave, appearing nearly flat (horizontal) in bud (petal axis nearly perpendicular to that of floral axis), not seen at anthesis, triangular, 5 – 6 × 8 mm, apex acute, base truncate-retuse, sides square, 1.5 mm thick, along the midline, outer surface densely covered in appressed sericeous golden hairs. Torus (raised surface of receptacle on which are inserted the stamens and carpels), not seen (material scarce). Stamens c. 150, spirally inserted, arranged in c.12 ranks. Stamens latrorse, basifixed, erect, linear 1 × 0.25 mm, filament minute, connective and thecae nearly the length of the stamen, connective terminal extension globose, 0.2 × 0.05 mm, black, forming an incomplete cap over the anther cells. Carpels 4, longer than stamens, subcylindrical, not dissected due to shortage of material. Stigma sessile, horseshoe shaped-crescentic, 0.8 × 0.6 mm, adaxial surface glabrous, outer surface with minute white patent hairs 0.05 mm long. *Fruit* monocarps (number per fruit unknown), drying brown, pericarp leathery, glossy, ellipsoid, 3.8 × 2.1 cm, not constricted between the seeds, apex rounded, base sessile, glabrous; Peduncle-pedicel, calyx-receptacle and bracts not recorded in fruit. *Seeds* c. 12 per monocarp, pale-yellow, oblong, compressed, widest at hilum end, 11 – 12 × 6 – 8 × 3 – 4 mm, marginal raphe ridge apex acute, hilum sunken, elliptic, 4 × 3 mm, margin raised, with short fibres, central sunken area with longitudinal ridge 2 mm long, micropylar aperture conspicuous; testa 0.2 mm thick, bony; endosperm ruminate in longitudinal section, embryo flat, c. 3 mm wide. Fig. 2 & 3.

##### RECOGNITION

*Lukea triciae* is similar to *Lukea quentinii* Cheek & Gosline, distinct in the leaves of reproductive stems being larger ((11.2 –)13.9 – 21 × (4.4 –)5.2 – 7.4(– 8.1) cm), oblong, lateral nerves 17 – 19 on each side, (versus smaller ((4.5 –)6.8 – 10.5 × (2.5 –)4 – 6 cm long), ovate or ovate-elliptic with lateral nerves 5 – 7(– 8) on each side of the midrib) and the larger flowers (petals 8 mm wide versus 5 – 6 mm wide), carpels 4 (versus (1–)2(–3)); fruits cylindric-ellipsoid, not constricted between the seeds (versus constricted between the seeds).

##### DISTRIBUTION

Tanzania (Map 1). Endemic to surviving lowland forest patches around the Udzungwa Mts

##### SPECIMENS EXAMINED. TANZANIA. Morogoro District

Mtibwa Forest Reserve, St. (Dec. 1953), *Semsei* 1520 (EA000008946, K000593315); Udzungwa Mts National Park, Msolwa - Camp 244, 7° 43’ S, 36° 55’ E, 350 m alt., fl. 23 Oct. 2005, *Luke WRQ, Mwangoka, Festo* 11205 (holo. K000593317); iso. EA000008947, MO, NHR); Magombera Forest 07° 48’S, 36°58’E, 250 m alt., fl. fr. 19 July 2003, *Q. Luke, P. Luke & Arafat* 9526 (EA, K000593316, MO3055171, NHT).

##### HABITAT

Evergreen lowland forest remnants; 250 – 350 m alt. At the Udzungwa Mts Msolwa location, *Lukea triciae* occurs on the bank of a small stream with riverine forest that runs through woodland at base of an escarpment. Associated species are *Uvaria tanzaniae, Pteleopsis myrtifolia, Terminalia sambesiaca, Sterculia quinqueloba, Pseudolachnostylis maprouneifolia, Erythrophleum suaveolens, Isoberlinia scheffleri, Parkia filicoidea, Tetrapleura tetraptera, Dalbergia* spp., *Celtis zenkeri, Ficus* spp., *Milicia excelsa, Treculia africana, Blighia unijugata, Lecaniodiscus fraxinifolius, Sorindeia madagascariensis, Pouteria alnifolia, Strychnos* spp., *Schrebera trichoclada, Diplorhynchus condylocarpon, Holarrhena pubescens, Catunaregam* spp., *Crossopteryx febrifuga, Borassus aethiopum*. At the Magombera Forest Reserve site, it was found in lowland ?groundwater forest with *Huberantha verdcourtii, Isolona heinsenii, Uvaria tanzaniae, Rinorea* spp., *Memecylon magnifoliatum, Cola* spp., *Rhodognaphalon schumannianum, Drypetes parvifolia, Erythrophleum suaveolens, Guibourtia schliebenii, Hymenaea verrucosa, Julbernardia magnistipulata, Albizia gummifera, Parkia filicoidea, Dalbergia* spp., *Pterocarpus mildbraedii, Ficus* spp., *Treculia africana, Pancovia holtzii, Sorindeia madagascariensis, Diospyros* spp., *Chionanthus mildbraedii, Aoranthe penduliflora, Coffea* spp., *Craterispermum schweinfurthii, Psychotria* spp., *Vitex doniana, Borassus aethiopum*.

##### CONSERVATION STATUS

The site of the earliest known collection of this genus is Mtibwa Forest Reserve (located at 6 07S, 37 39E according to Polhill 1998). According to https://www.tfs.go.tz/index.php/en/forests/mtibwa (accessed 21 April 2021), it is now a forestry plantation, 95 % being teak, *Tectona grandis* L.f., and only a few small scraps of indigenous vegetation remain on land unsuitable for plantation. This can be confirmed by viewing through Google Earth using the georeference provided above where the former forest is now divided into parcels of planted trees. It seems likely, even certain, that the species has become extinct at this location since it was collected by Semsei in 1953. The species is therefore now confined to two sites separated from each other by 12 km of cultivated land at the foot of the Udzungwa Mts on current evidence. Since there are only two collections, one from each site, we estimate the maximum area of occupation as being 8 km^2^ using the preferred gridcells of 4 km^2^ favoured by IUCN (2012).

The Magombera Forest Reserve site where the species was discovered in 2003, was highly threatened when it was degazetted from the Selous Game Reserve (now also known as the Nyerere National Park) and not regazetted immediately as was intended at the time. It was part privately owned by the adjacent sugar estate that cleared much of the previous forest in the adjoining Kilombero valley in the 1970s. (https://www.rainforesttrust.org/projects/creating-the-magombera-nature-reserve-in-tanzania/ Accessed 8 May 2021). The Tanzania-Zambia railway was also cut through the forest in the 1970s.

A recent campaign to have Magombera given special protection has been successful. It was gazetted as a Nature Reserve in 2019 and is now subject to fundraising. Since 2008 (imagery viewed via Google Earth) some patches of forest at the margin have been cleared, presenting a threat to the habitat of *Lukea triciae*. Threats at the Msolwa site are firewood collection (Luke pers. obs.). Given the threats above and small AOO, we here assess the species as EN B2ab(iii).

##### PHENOLOGY

flowers and fruit in July, flower buds in Oct., sterile in December.

##### ETYMOLOGY

Named for the co-collector (with Quentin Luke) of material of this species, Patricia (Tricia) Luke.

##### VERNACULAR NAME & USES

None are recorded.

##### NOTES

*Lukea triciae* is more incompletely known than *L. quentinii*. Only a single monocarp is known, and the few flowers available are not quite at anthesis. The earliest, gathering, (*Semsei* 1520, K, sterile) has leaves of similar size and shape to those of fertile collections and so there is no evidence that there is a different juvenile or non-flowering stage, as seen in *L. quentinii* (see above). *Lukea triciae* is confined to Morogoro District, Tanzania, to which numerous other rare and threatened species are globally endemic, e.g. *Brachystephanus schleibenii* (Mildbr.) Champl., (Champluvier & Darbyshire 2009), and *Keetia semsei* Cheek & Bridson (Cheek & Bridson 2019) both known from a single collection, and *Asystasia asystasioides* Darbysh. & Ensermu (Darbyshire & Kelbessa 2007).

## Discussion

### The Eastern Arc Mountains & Coastal Forests

The two species are separated geographically by c. 525 km, as measured by Google Earth. Both locations are placed in the Eastern Arc mountains and Coastal Forests (EACF). The EACF are found in Tanzania and southern Kenya. They form an archipelago-like phytogeographical unit well-known for high levels of species endemism in many groups of organisms (Gereau *et al*. 2016). Among the better-known mountain blocks are the Udzungwas, the Ulugurus, and the Usambaras. The Mtibwa Forest reserve at the foot of the Udzungwas, home of *Lukea triciae*, is among the least known, least well surveyed, and least protected forest patches. Supported by moist air currents from the Indian Ocean, the surviving evergreen forests of the Eastern Arc Mountains alone have 223 species of endemic tree (Lovett 1998) and are variously stated to have 800 (Tanzanian Forest Conservation Group, undated) or as many as 1500 species (Skarbek 2008) of endemic plant species. In herbaceous groups such as the Gesneriaceae, over 50% of the species for East Africa (Uganda, Kenya and Tanzania) are endemic to the Eastern Arc Mts (Darbyshire 2006) and in the Acanthaceae, there are as many as 100 species endemic to the Eastern Arc Mts (Darbyshire *et al*. 2010) e.g. *Isoglossa bondwaensis* I. Darbysh. endemic to the Uluguru Mts (Darbyshire 2009). In terms of documented plant species diversity per degree square, the Eastern Arc Mts are second only in tropical Africa to S.W. Cameroon in the Cross-Sanaga Interval of W-C Africa (Barthlott *et al*. 1996, Cheek *et al*. 2001), which also has the degree square with highest generic diversity (Dagallier *et al*. 2020). *Lukea* is an example of a genus endemic to the EACF, just as e.g. *Medusandra* Brenan (Peridiscaceae, Soltis *et al*. 2007; Breteler *et al*. 2015) is endemic to the Cross-Sanaga interval. Several forest genera have disjunct distributions, being found only in the Cross-Sanaga Interval and in the EACF and not in between, e.g., *Zenkerella* Taub. and *Kupea* Cheek (Cheek *et al*. 2003, Cheek 2004b). The EACF include the sole representatives of plant groups otherwise restricted on the continent to the forests of Guinea-Congolian Africa, e.g., *Afrothismia* Schltr. and *Ancistrocladus* Wall. (Cheek & Jannerup 2006, Cheek *et al*. 2000). Extensive forest clearance within the last 100 – 150 years has removed forest from some mountains entirely, and reduced forest extent greatly in others. Since the 1970s more than 12% of these forests have been cleared (Tanzania Forest Conservation Group, undated). Yet new taxa such as these two *Lukea* species are still being published steadily. Let us hope that their natural sites where not already destroyed can be preserved and their extinction avoided.

### Conclusions

*Lukea quentinii* and *L. triciae* underline the urgency for identifying and publishing species while they still survive. Threats to such new discoveries for science are ever present, putting these species at high risk of extinction. About 2000 new species of flowering plant have been published by science each year for the last decade or more (Cheek *et al*. 2020), adding to the estimated 369 000 already documented (Nic Lughadha *et al*. 2016), although the number of flowering plant species known to science is uncertain (Nic Lughadha *et al*. 2017). Only 7.2% of species have been assessed and included on the Red List using the IUCN (2012) standard (Bachman *et al*. 2019), but this number rises to 21 – 26% when additional evidence-based assessments are considered, and 30 – 44% of these assess the species as threatened (Bachman *et al*. 2018). Newly discovered species, such as that reported in this paper, are likely to be threatened, since widespread species tend to have been already discovered. There are notable exceptions to this rule (e.g. *Vepris occidentalis* Cheek (Cheek *et al*. 2019) a species widespread in West Africa from Guinea to Ghana). However, it is generally the more localised, rarer species that remain unknown to science. This makes it all the more urgent to find, document and protect such species before they become extinct. Until species are delimited, described and known to science, it is difficult to assess them for their IUCN conservation status and so the possibility of protecting them is reduced (Cheek *et al*. 2020). Documented extinctions of plant species are increasing, e.g., in coastal forest of Kenya *Cynometra longipedicellata* Harms may well now be extinct at its sole locality, the Amani-Sigi Nature Reserve (Gereau *et al*. 2020) and in Tanzania *Kihansia lovettii* Cheek (Cheek 2004b) at the Kihansi dam site, has not been seen since it was first collected despite targeted searches. There are also examples of species that appear to have become extinct even before they are known to science, such as *Pseudohydrosme bogneri* in Gabon (Moxon-Holt & Cheek 2021) and in Cameroon *Vepris bali* Cheek (Cheek *et al*. 2018b). In all cases anthropogenic pressures have been the cause of these extinctions. In both Kenya and Tanzania, on the whole, natural habitat is relatively well covered by a well-planned network of protected areas, but nevertheless natural habitat at some sites with species of high value for conservation has all but disappeared as for the site for first collection of *Lukea triciae* at Mtwiba (see above). Further investment in prioritising the highest priority areas for plant conservation as Tropical Important Plant Areas (TIPAs, using the revised IPA criteria set out in Darbyshire *et al*. (2017)) as is in progress in countries such as Guinea, Cameroon, Mozambique and Ethiopia might be extended elsewhere to reduce further the risk of future global extinctions of range-restricted endemic species such as *Lukea quentinii* and *L. triciae*.

## Acknowledgements

We thank Dr Kaj Vollesen for making available to us the material published in this paper as *Lukea*, which he had been first to identify as a new genus. We thank Nina Davies for facilitating this. Janis Shillito typed the manuscript. Thomas Couvreur and Leo-Paul Dagallier of Univ Montpellier are thanked for accelerating completion of this paper and for generously sharing the unpublished results of their molecular phylogenetic research on the *Uvariopsis* clade.

